## Supplemental Information, figures with legends for "Sex-specific attenuation of photoreceptor degeneration by reserpine in a rhodopsin P23H rat model of autosomal dominant retinitis pigmentosa"

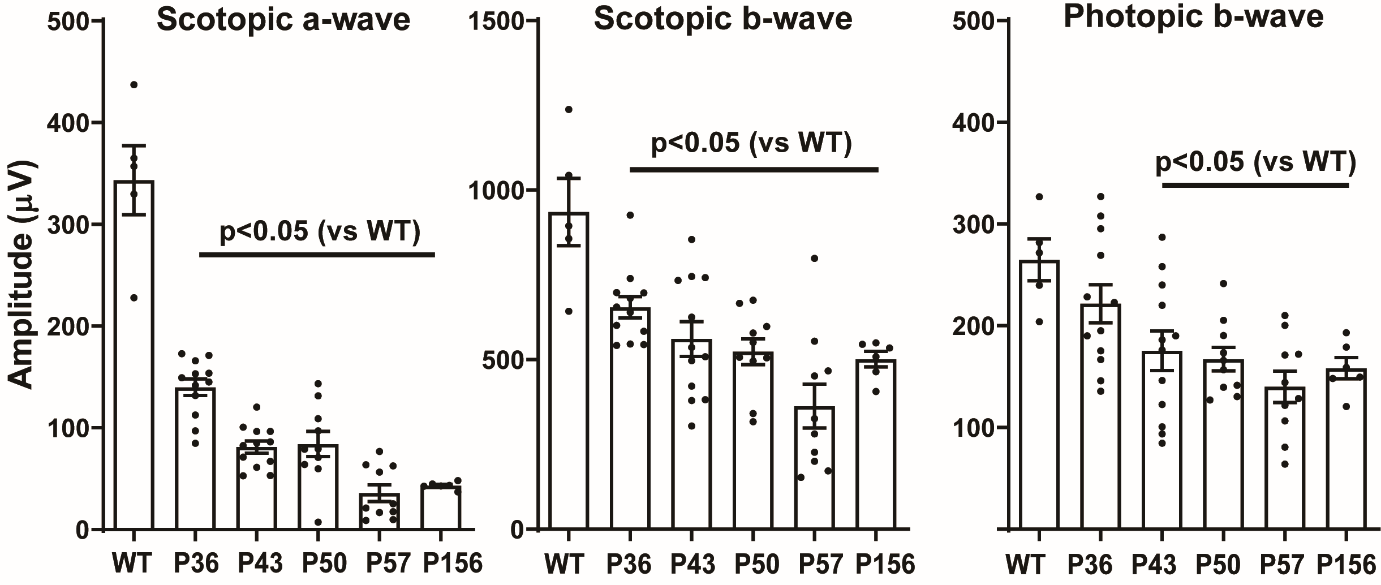

**Supplementary Figure 1. Progression of retinal degeneration in P23H-1 rats evaluated by electroretinography.**

**Scotopic and photopic electroretinogram responses at P36, P43, P50, P57, and P156 in P23H rats and at P78 in wild type rats.** All parameters were measured in both eyes (3 to 6 P23H-1 rats and 3 wild type rats). Data are expressed as mean ± SEM, and the Mann-Whitney test is used to compare P23H-1 and wild type rats. WT: wild type.

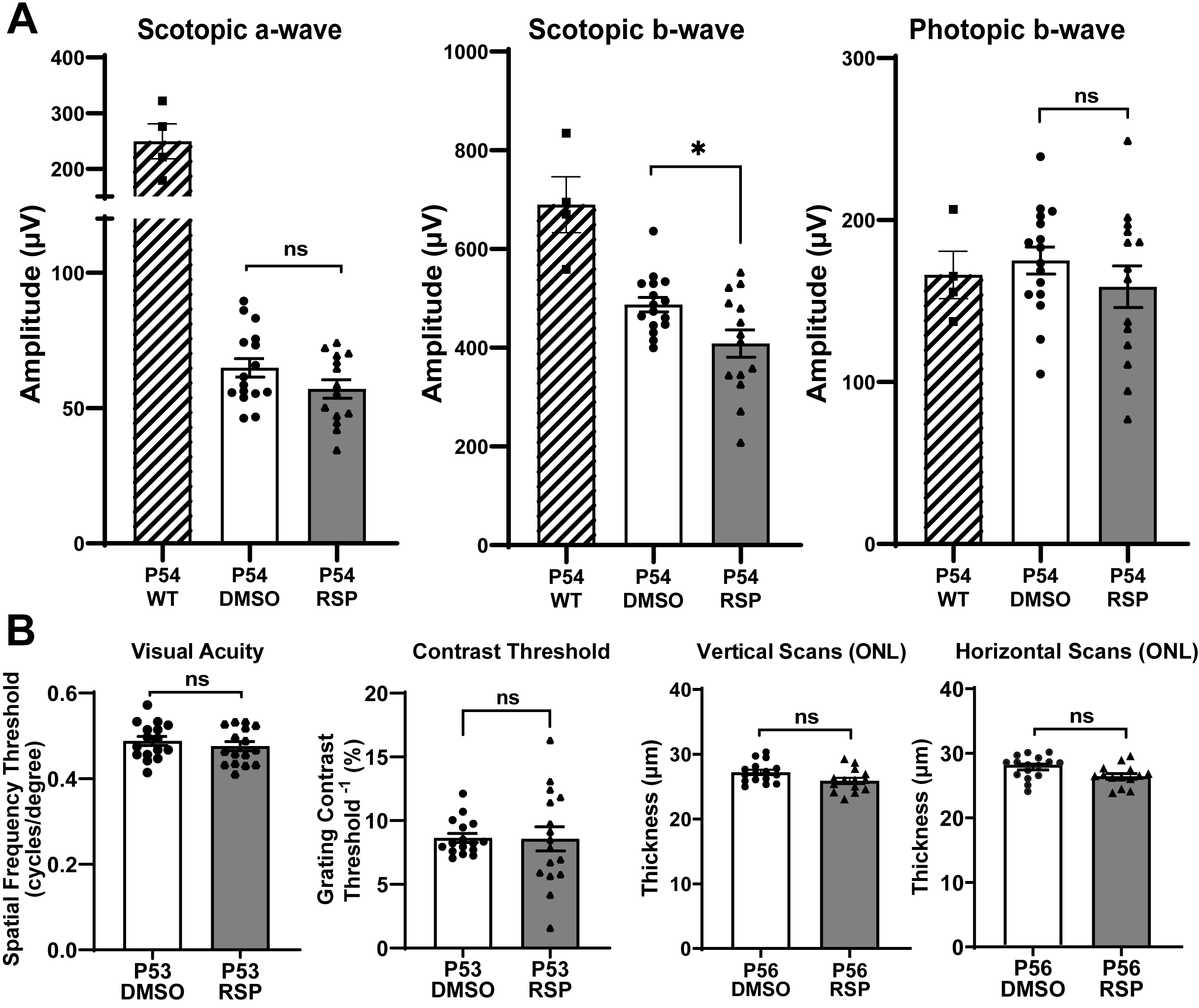

**Supplementary Figure 2. Treatment responses in P23H-1 rats.**

**(A) Scotopic and photopic electroretinogram responses at P54** in wild type, DMSO- and RSP-treated P23H-rats. **(B)** Visual acuity measured by OKT response and contrast threshold at P53, and outer nuclear layer thickness measured by vertical and horizontal scan of OCT at P56 in DMSO- and RSP-treated P23H-rats. All parameters were measured in both eyes (2 wild type, 8 DMSO-treated and 7 RSP-treated rats). Data are expressed as mean ± SEM, and the Mann-Whitney U test is used to compare DMSO- and RSP-treated groups. RSP: reserpine, OKT: Optokinetic tracking, OCT: Optical coherence tomography, ONL: outer nuclear layer, ns: not significant, *p<0.05.

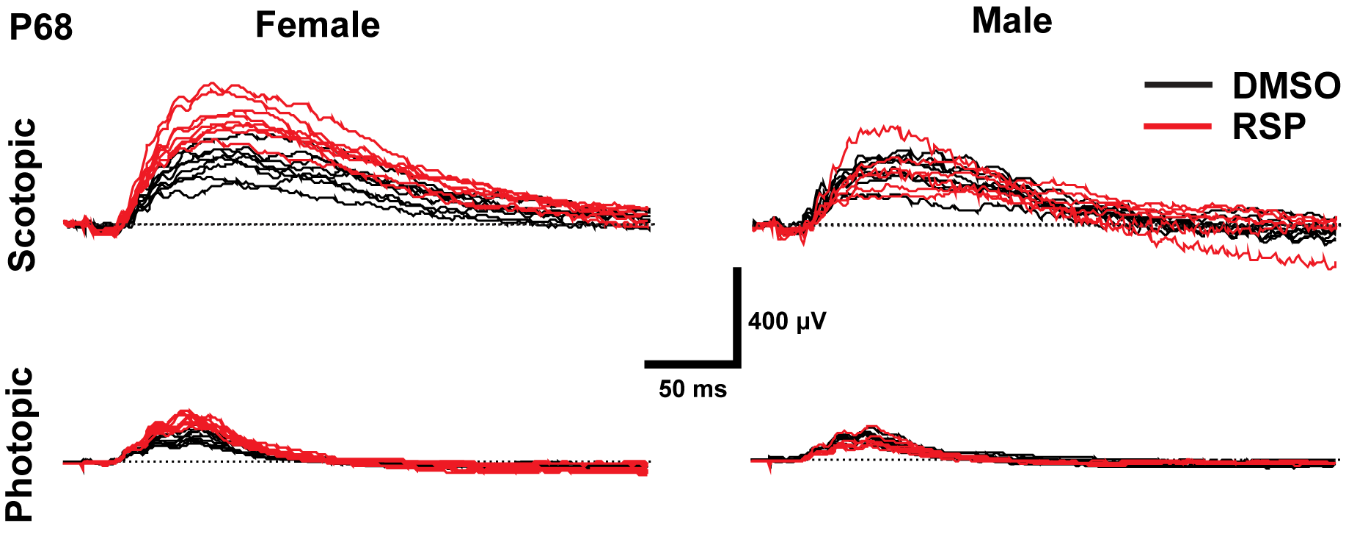

**Supplementary Figure 3. ERG responses in P23H-1 rats grouped by sex.**

Scotopic and photopic electroretinogram responses at P68 in P23H rats. Responses were measured in both eyes (4 DMSO-treated and 4 RSP-treated female rats and 4 DMSO-treated and 3 RSP-treated male rats). RSP: reserpine.

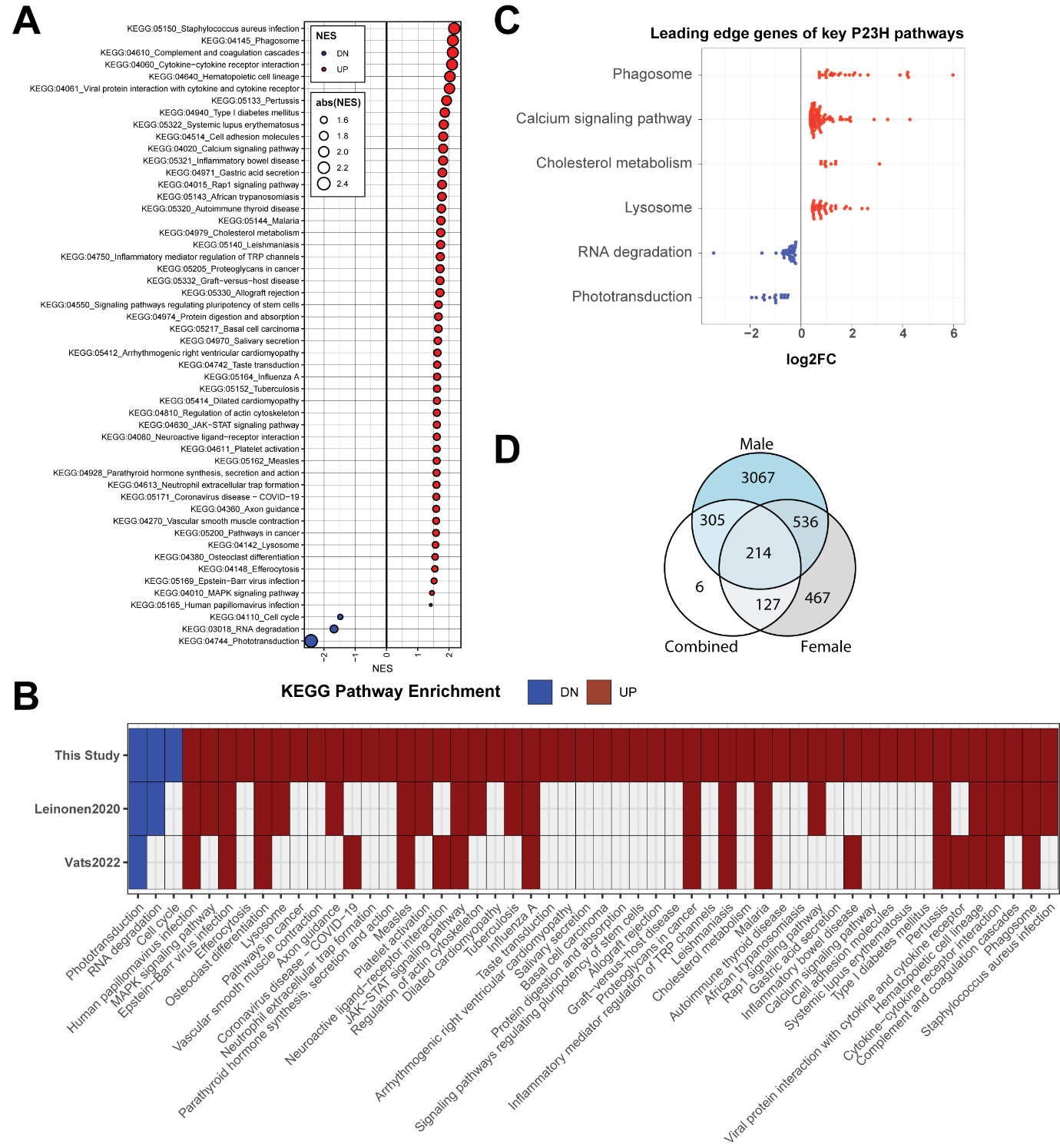

**Supplementary Figure 4. Rho P23H linked molecular pathology is consistent across different model systems.**

(**A**) Summary of significantly enriched KEGG processes in P23H mutants when compared to WT. Female and male samples were combined in this analysis. (**B**) Pathways enriched in P23H retinal transcriptomes, when compared to WT, are consistent across model systems. Qualitative heatmap shows up- and down- regulated KEGG pathways in our study as well as published (Leinonen et al., 2020^[1]^; Vats et al., 2022^[2]^). Pathways are colored red and blue per over- and under-enrichment, and grey if not significant in a dataset. (**C**) Beeswarm plot of log2 fold change of leading-edge genes for selected pathways, showing increase in phagosome as well as lysosome processes and reduction in phototransduction. (**D**) Female and male retinas have distinct sets of P23H linked differential expression. Venn diagram comparing differentially expressed genes from combined samples, and from independent male and female comparisons.

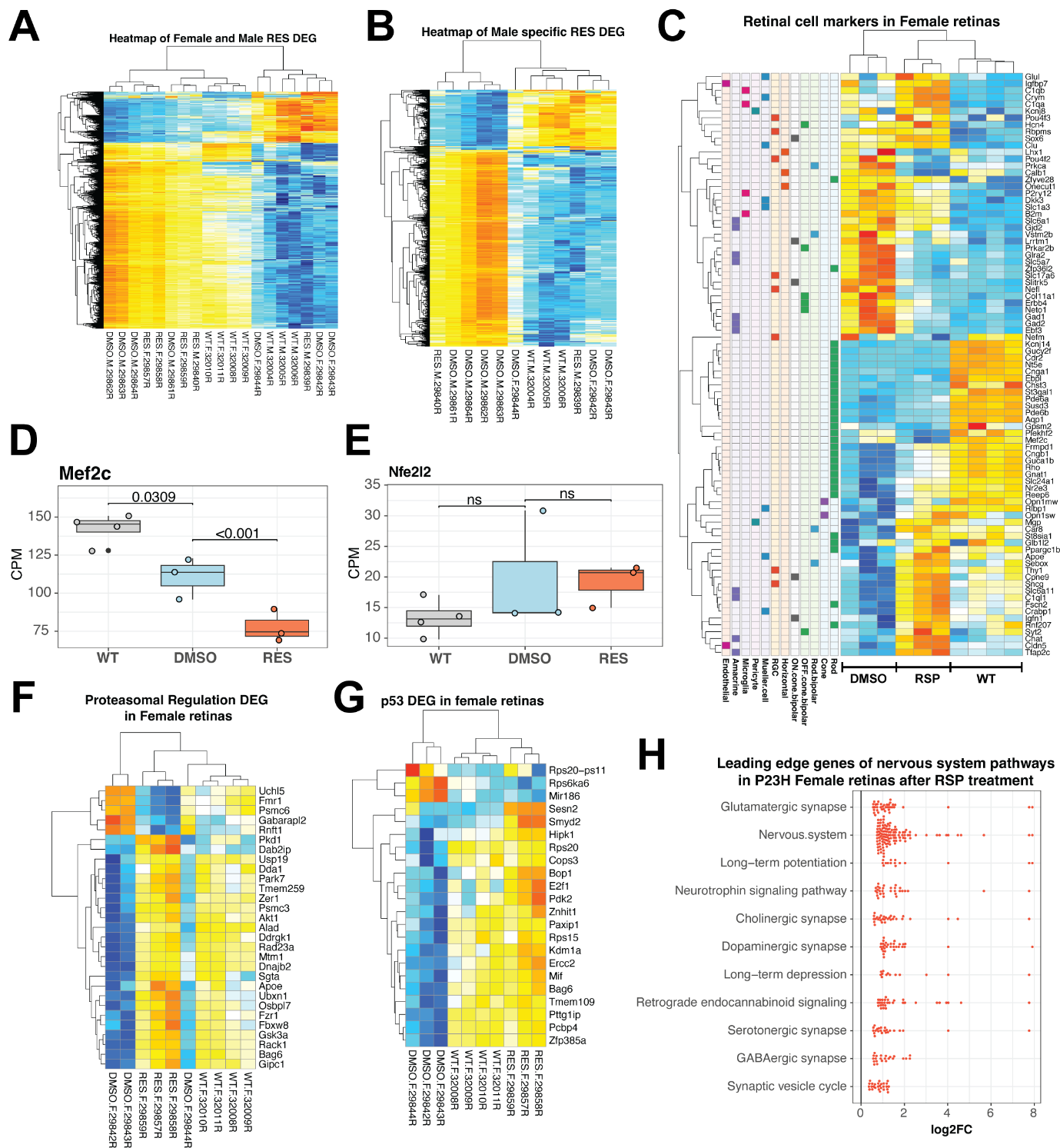

**Supplementary Figure 5. Distinct transcriptomic response to reserpine in female and male mutant retinas.**

Heatmaps of differential genes from (**A**) combined female and male, and (**B**) male only analysis. Colors are z-scores of row-scaled log2 CPM values of genes and same as in Figure 5B. (**C**) Gene expression profile of retinal cell type markers show reserpine linked transcriptomic changes in rod photoreceptors as well as other cell types. Colors are z-scores of row-scaled log2 CPM values of genes and same as in Figure 5B. (**D**) Boxplots of gene expression of retinal transcription factor Mef2c in WT, untreated and reserpine treated female retinas. (**E**) Boxplots of expression of the gene coding for proteasomal regulator Nfe2l2 (Nrf2) in WT, and RSP treated and untreated P23H female retinas. Heatmap of significantly differential genes in (**F**) proteasomal regulation and (**G**) p53 signaling pathways in female retinas. Colors are z-scores of row-scaled log2 CPM values of genes and same as in Figure 5B. Adjusted p-value from statistical comparison is shown on the plot, with “ns” denoting a value greater than 0.05. (**H**) Plot showing gene expression alteration (log2 fold change) among leading edge genes for KEGG nervous system gene sets. Term “Nervous.system” is a merged superset for all sub-pathways in that category. WT: wild type, DMSO: DMSO treated P23H retinas, RSP: reserpine treated P23H retinas.

**Supplemental Table Legends**

**Table S1.** Differential gene expression between WT and P23H retinas.

**Table S2.** Differential gene expression in P23H retinas after reserpine treatment.

**Table S3.** Individual values of ERG and optokinetic tracking responses in DMSO- and RSP-treated rats. RSP = reserpine.

1. Leinonen, H., Pham, N. C., Boyd, T., Santoso, J., Palczewski, K., & Vinberg, F. (2020). Homeostatic plasticity in the retina is associated with maintenance of night vision during retinal degenerative disease. *Elife*, *9*. https://doi.org/10.7554/eLife.59422
2. Vats, A., Xi, Y., Feng, B., Clinger, O. D., St Leger, A. J., Liu, X., Ghosh, A., Dermond, C. D., Lathrop, K. L., Tochtrop, G. P., Picaud, S., & Chen, Y. (2022). Nonretinoid chaperones improve rhodopsin homeostasis in a mouse model of retinitis pigmentosa. *JCI Insight*, *7*(10). https://doi.org/10.1172/jci.insight.153717
