## Supplementary figures and images for "Sex-specific attenuation of photoreceptor degeneration by reserpine in a rhodopsin P23H rat model of autosomal dominant retinitis pigmentosa"

### Appendix 1 - Figure 1

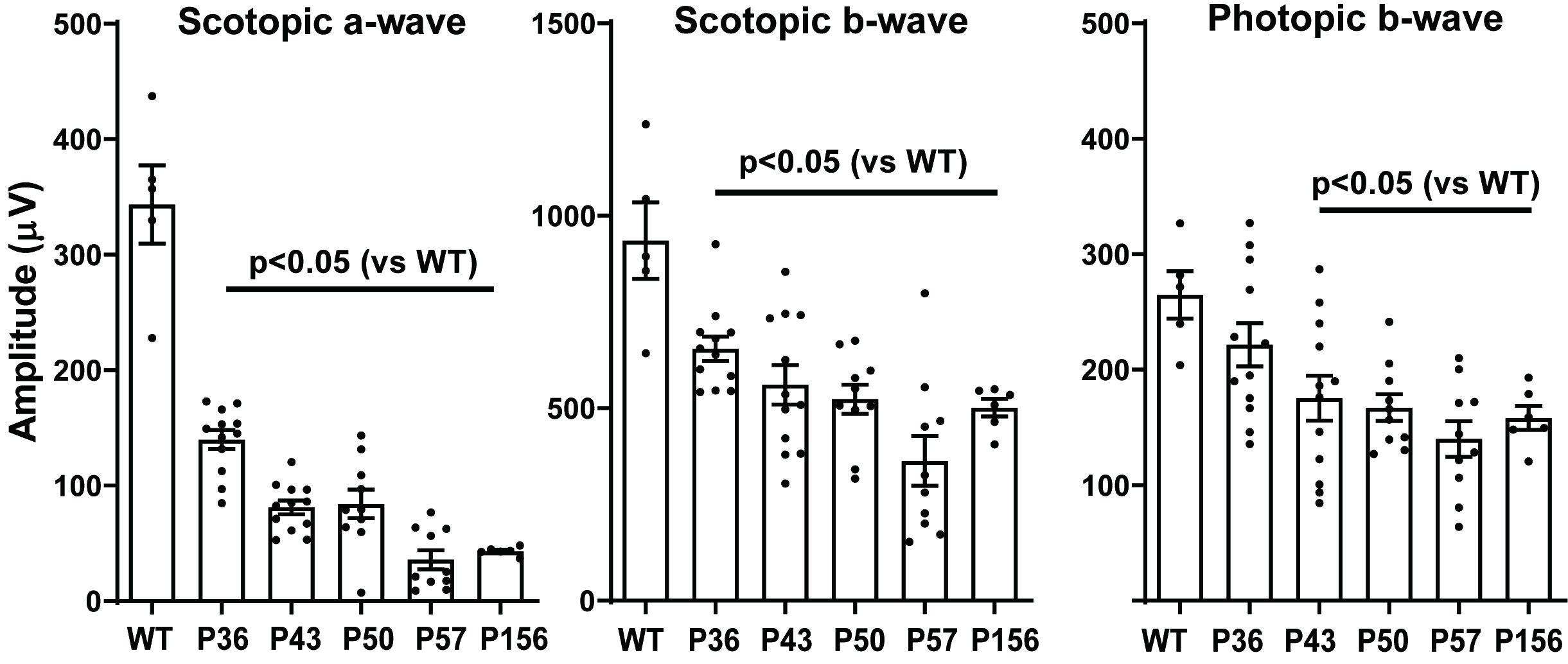

### Figure 1 - Figure Supplement 1

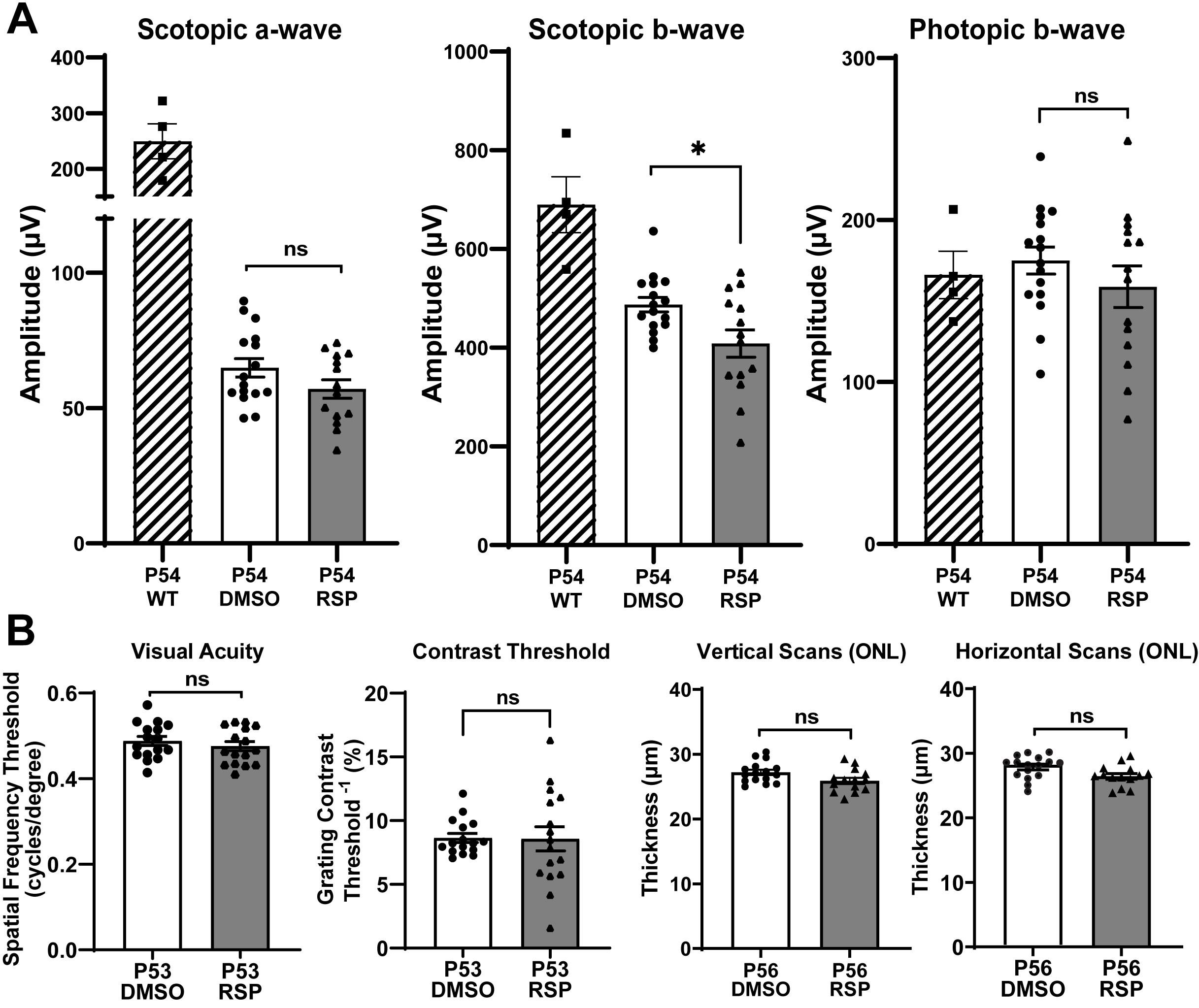

### Figure 2 - Figure Supplement 1

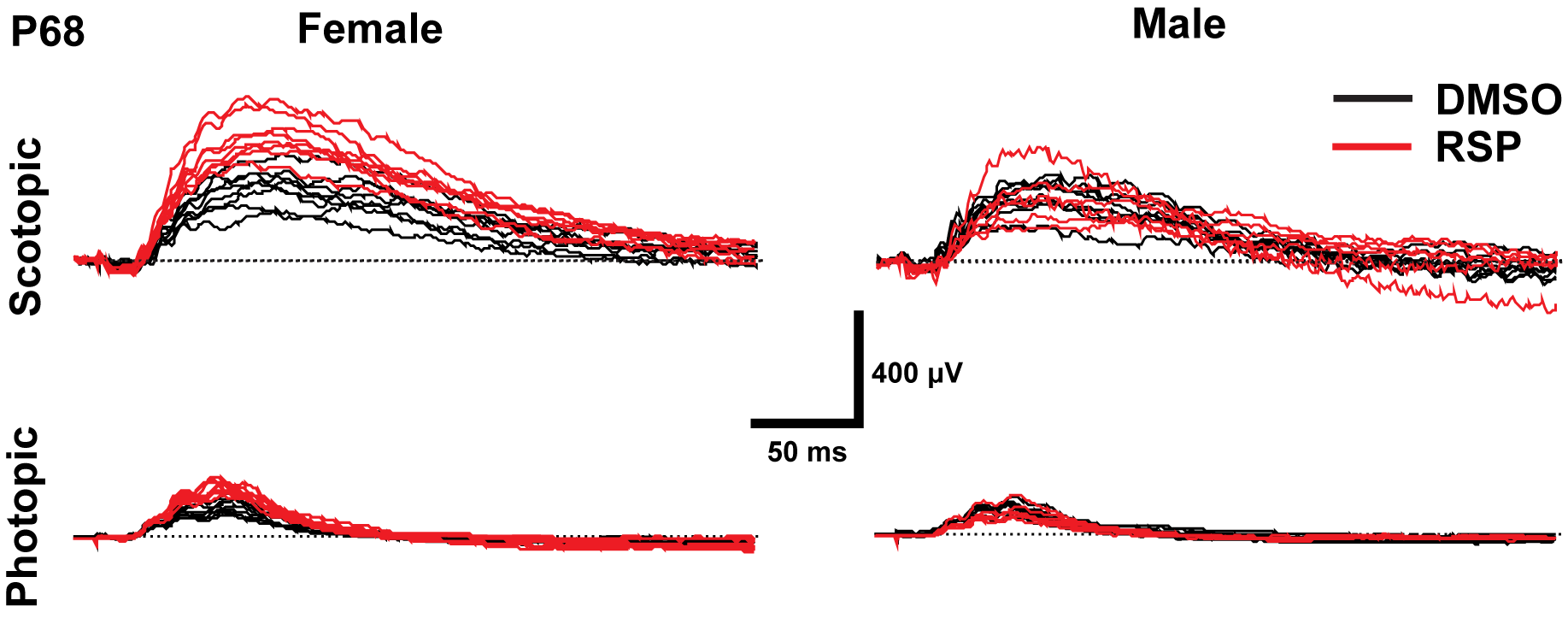

### Figure 5- Figure Supplement 1

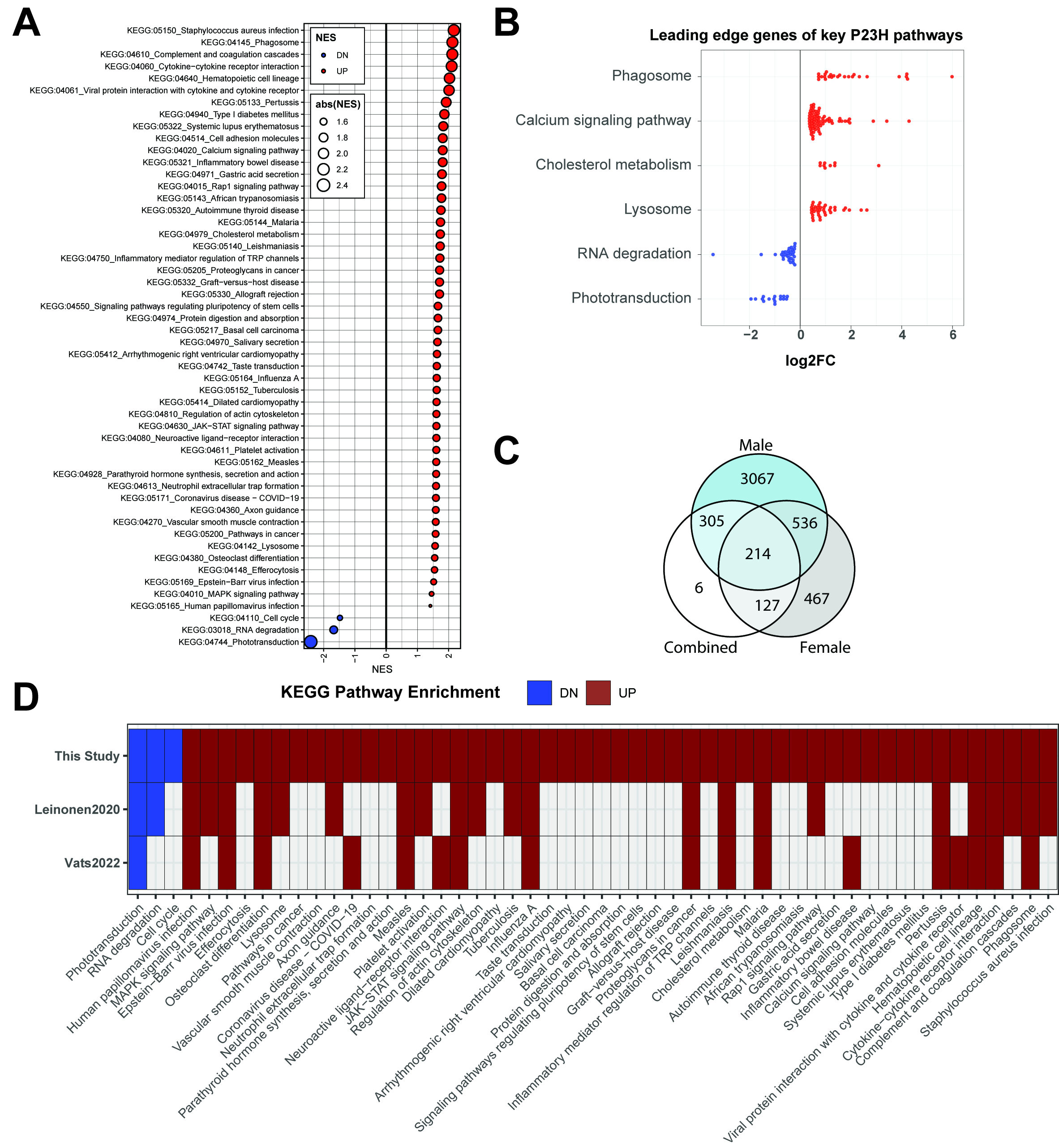

### Figure 6 - Figure Supplement 1

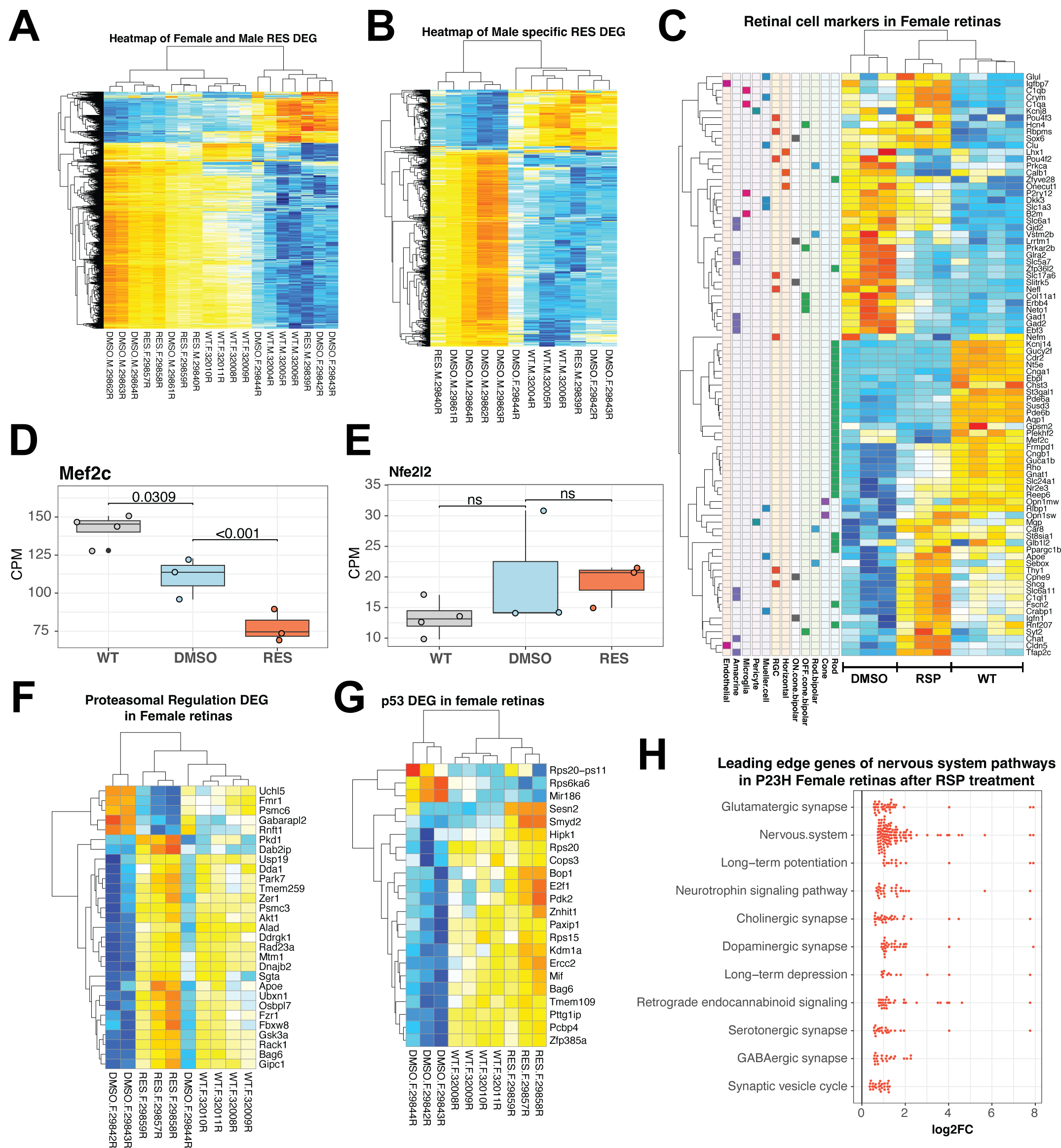
